# A hybrid framework integrating structural machine learning and 3D liver-on-chip assay for drug-induced liver injury prediction

**DOI:** 10.64898/2026.06.15.732231

**Authors:** Fan Zhang, Yu Zhou, Duanchen Ding, Feng Zhang, Rong-Rong Xiao, Xiaoni Ai

## Abstract

Drug-induced liver injury (DILI) remains a major cause of clinical attrition and postmarketing withdrawal, but structure only DILI predictors are difficult to compare because public benchmarks are vulnerable to compound overlap, scaffold similarity and shared label provenance. We present **Oakuloid^TM^**, an open DILI prediction framework that pairs a leakage audited structure based model with an optional iBAC 3D primary human hepatocyte IC**_50_**/C**_max_** confirmation signal. The structural model integrates gradient boosted descriptor backbones, fingerprint random forests and LivTox proxy-DILI features through a logistic meta learner. Its evaluation is designed as part of the contribution: internal DILIrank, strict external TDC, scaffold disjoint TDC and independent Geci provenance checks are reported with released per compound predictions. Oakuloid reaches AUROC **0.811** on the strict external TDC benchmark and remains competitive under scaffold and fully clean TDC filtering. A channel attribution ablation shows that the external benchmark lead is driven by descriptor based gradient boosted trees rather than by DILIPredictor derived proxy features, reducing a potential circularity concern. The wet lab IC**_50_**/C**_max_** signal is largely orthogonal to structure and supports a confirmation mode that shifts the internal operating point toward higher specificity without claiming a universal AUROC gain. Oakuloid is released with code, model artifacts, calibration analysis, a 122 compound wet lab benchmark and a model card under the Apache License 2.0, supporting reproducible DILI screening and benchmark auditing.

## 1 Introduction

Drug-induced liver injury (DILI) accounts for approximately half of all acute liver failure cases in industrialised countries [63, 69] and contributes to 25–30% of late stage drug development attrition [9, 12, 31, 48, 66, 70]. Predicting clinical hepatotoxicity at the candidate selection stage therefore remains a central, unresolved challenge in computational drug safety.

Two complementary classes of DILI predictor have emerged, and both face distinct performance limits. Structure only methods are computationally inexpensive, deterministic, and applicable to any compound with a SMILES string, but their public benchmarks have plateaued at an area under the receiver operating characteristic curve (AUROC) of approximately 0.75 over the past decade [41, 55, 72], a range that has persisted despite extensive architectural exploration [1, 15, 20, 34, 40, 46, 54, 55, 59, 75], suggesting the bottleneck is the signal class itself: a single SMILES carries insufficient information to resolve the mechanistically heterogeneous causes of hepatotoxicity [29–31, 62]. Recent deep models report markedly higher figures, with AUROC 0.88–0.97 on random or stratified splits of shared-provenance reference sets [37, 38], but such gains largely track train–test similarity rather than generalization: DILIGeNN itself reports an established predictor falling to AUROC ≈ 0.63 under a scaffold split [38], near-duplicate train–test molecules are a documented source of inflated molecular-ML performance [58], and a 2025 pretrained graph model evaluated without such overlap remains at AUROC ≈ 0.76 [24], consistent with the observed structural performance range. Public predictors are also rarely released with multiseed variance reporting or strict external leakage audit [23, 32, 60, 67], making method to method comparisons difficult; and most learn from structure alone, even though exposure-normalised cellular cytotoxicity (the ratio of half-maximal inhibitory concentration to maximum plasma concentration, IC_50_/C_max_) is an established orthogonal DILI risk signal [3, 13, 56, 61]. Treating public clinical reference lists as purely structural tasks can therefore omit a signal already used in drug safety practice.

Wet lab DILI screens are equally constrained. HepG2 high content cytotoxicity reaches concordance of ∼50–65% on canonical reference sets [47, 74]; conventional 2D primary human hepatocyte (PHH) cytotoxicity assays reach AUROC ∼0.70 and lose cytochrome P450 (CYP450) activity within 48–72 h [6, 64], underdetecting metabolite-mediated DILI; higher fidelity 3D microphysiological systems, including recent human liver organoid and liver-on-chip platforms, improve this to AUROC ∼0.75 [5, 7, 27, 35, 39, 43, 49, 51, 78] but at an order of magnitude higher per compound cost in reagents, procurement, and assay time, and a single 72-hour cellular endpoint cannot register Type-B (idiosyncratic, immune-mediated, or chronic exposure) DILI [9, 31]. The structural and wet lab signals are therefore *complementary rather than substitutable*, but they have rarely been paired in a single open and auditable pipeline.

To address these complementary limitations, we present **Oakuloid**, a hybrid framework that pairs an enhanced structure only classifier with a wet lab fusion module. The structural component is a stacked ensemble whose three gradient boosted decision tree (GBDT) backbones (LightGBM, XGBoost, CatBoost), four fingerprint random forests (MACCS, Avalon, atom-pair, topological-torsion), and four predicted LivTox proxy-DILI features are combined by a logistic meta learner. CatBoost hyperparameters are selected by groupwise (InChIKey-14) cross validated Optuna search on the training set alone, while LightGBM, XGBoost and fingerprint random forests use prespecified class balanced configurations; no external or internal evaluation label or outcome is used for tuning or model selection. The fusion module combines this structural score with the experimental IC_50_/C_max_ ratio by rank mean averaging. The ratio is measured on our previously reported 3D-PHH *integrated biomimetic array chip* (iBAC) platform [73], adding a measured cellular damage signal that is not directly observable from structure. The full pipeline is released under the Apache License 2.0 (Figure 1).

**Fig. 1.**
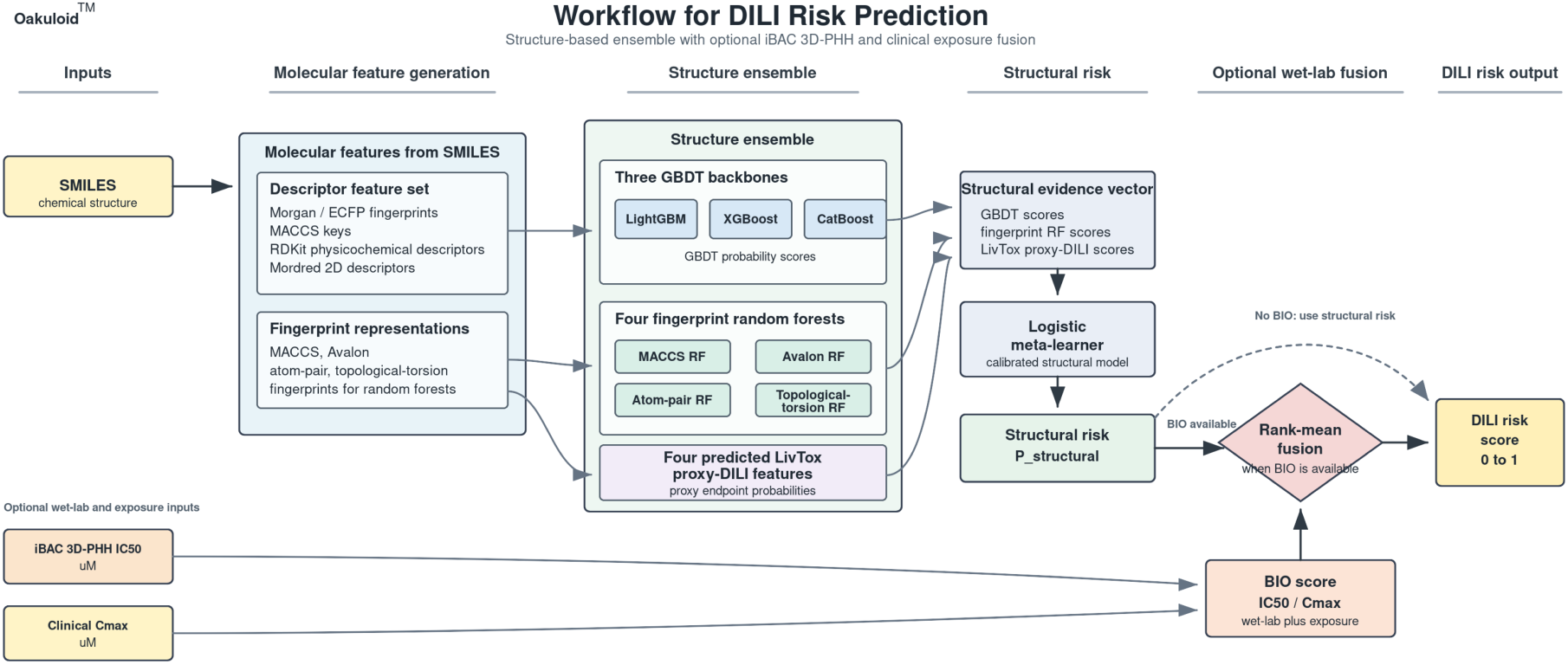
Oakuloid workflow. SMILES strings are converted to standard molecular descriptors, routed through three GBDT backbones, four fingerprint random forests and four predicted LivTox proxy-DILI features, and combined by a logistic meta learner. When an experimental iBAC IC_50_/C_max_ measurement is available, the structural score and wet lab ratio are combined by the rank mean fusion rule; otherwise the structural score is used directly.

This work makes three contributions:

1. **A leakage audited, multi-axis DILI benchmark with an open release.** Oakuloid is evaluated across standard, scaffold disjoint and label-provenance-disjoint settings that separate ordinary benchmark performance from novel-chemistry generalization; the model code, inference pipeline, 122-compound DILIrank wet lab benchmark, per-compound predictions, calibration analysis and a Mitchell style model card [44] are released under Apache License 2.0 to support reproducible comparison.
2. **A structural stack whose lead is attributed, not assumed.** Oakuloid leads the out-of-distribution-valid baselines on the standard and scaffold disjoint benchmarks, and a channel-attribution ablation shows this lead comes from the gradient boosted descriptor backbones rather than from the DILIPredictor-derived proxy features, reducing the circularity concern in comparisons with DILIPredictor.
3. **An optional wet-lab confirmation mode with explicit tradeoffs.** The iBAC 3D-PHH IC_50_/C_max_ ratio carries information not recovered from structure alone (*R*^2^ ≈ 0); a single rank mean rule converts a high-sensitivity structural screen into a higher-specificity confirmation step, with the sensitivity/specificity tradeoff reported in full.

## 2 Results

### 2.1 Model architecture and external benchmark performance

Oakuloid’s default structure-only predictor is the four proxy stack shown in Figure 1. A SMILES string is first converted to a shared molecular descriptor matrix and to four complementary fingerprint representations. The descriptor matrix is scored by three gradient boosted decision-tree backbones (CatBoost, LightGBM and XGBoost), while the fingerprint views are scored by four random forest models. Four predicted LivTox proxy-DILI endpoints are then appended as additional structure-derived evidence channels. These eleven inputs are combined by an L2-regularized logistic meta learner to produce the shipped structural probability score.

We use this smaller four proxy stack as the primary model because it keeps the deployed feature vector compact and makes the channel attribution analysis transparent. A larger ten proxy panel is retained only as a robustness ablation in the supplement, not as the primary deployed model. When an experimental iBAC 3D-PHH IC_50_/C_max_ ratio is available, it is not folded into the structural model; instead, the structural probability and the wet lab ratio are combined downstream by the rank mean fusion rule described in Section 5.7. Thus the structure-only screen and the optional wet lab confirmation mode remain separable.

#### External benchmark performance across datasets

We first report structure-only AUROC on three evaluation sets before interpreting stricter leakage-audit subsets in Section 2.2. The strict external TDC subset is the primary public benchmark; the internal DILIrank *N* = 94 cohort is shown here only as a structure-only reference because the same cohort is used later for wet lab fusion; and the Geci independent set tests transfer to a different label provenance. Figure 2 therefore summarizes ordinary model ranking across the three datasets, while the scaffold-disjoint, fully clean and doubly disjoint analyses are handled separately as leakage audits.

**Fig. 2.**
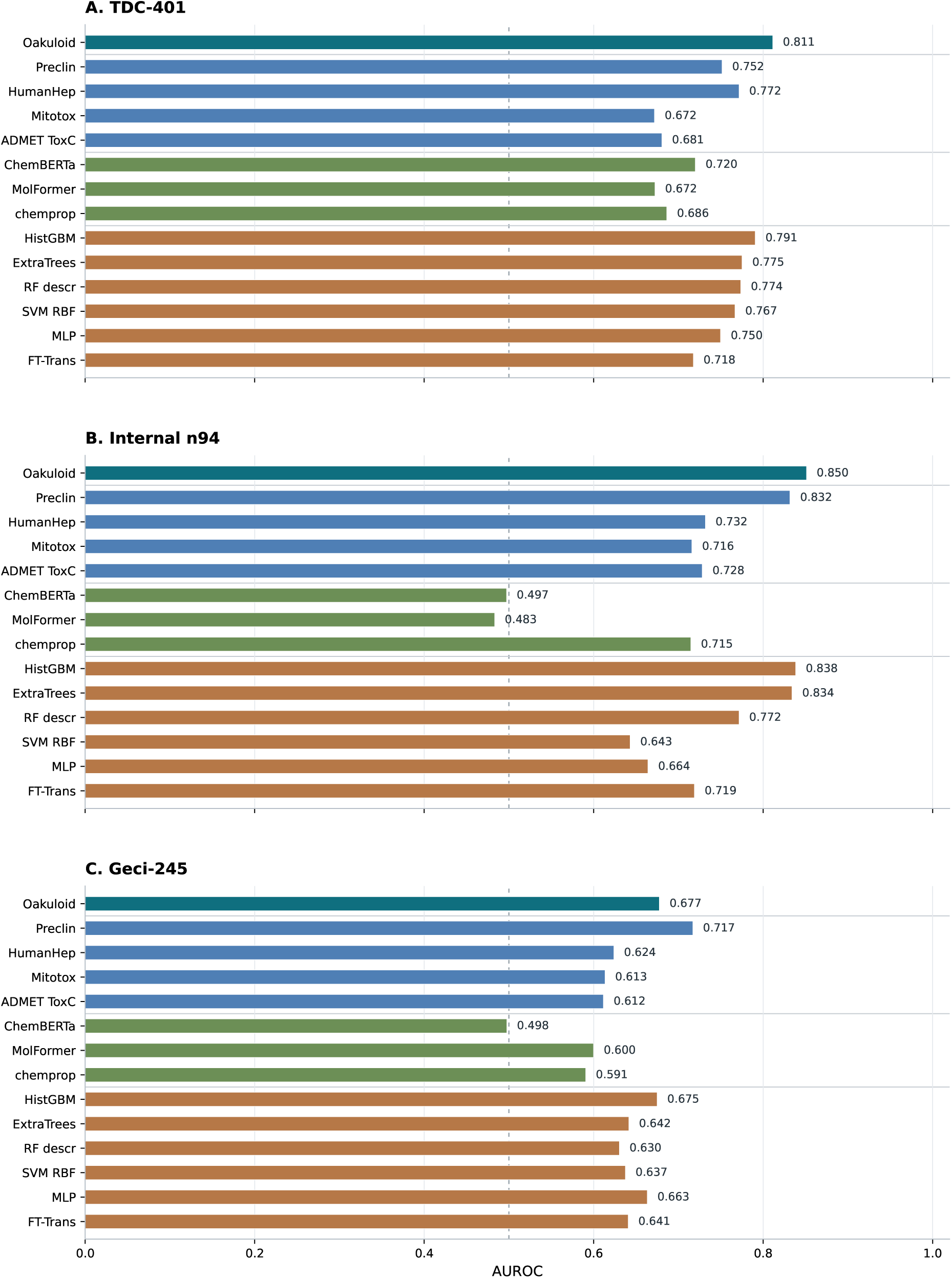
An external model comparison. AUROC on three evaluation sets. Dashed line marks random ranking.

On the strict external TDC benchmark (*n* = 401, 196 positive / 205 negative), Oakuloid reaches AUROC **0**.**811** (95% CI 0.77–0.86; five seed mean 0.809 ± 0.001; Figure 2A). Its AUROC is significantly higher than the principal public baselines by DeLong’s test (DILIPred-Preclin [55], ChemBERTa-2 [1], chemprop [76] and MolFormer-XL [54]; all *p <* 10*^−^*^3^). The +0.06 AUROC lead over the principal public comparator, DILIPred-Preclin, is stable across training seeds and is positive in all five, per-seed +0.061 ± 0.001. Two further DILIPredictor endpoints (DILIPred-HumanHep, DILIPred-Mitotox) and an ADMET-AI toxicity-consensus head are included in the same comparison (Figure 2). On the internal DILIrank structure-only reference set, Oakuloid reaches AUROC 0.850, again ahead of the evaluated structure-only alternatives (Figure 2B). On the Geci independent set, however, DILIPred-Preclin is numerically higher than Oakuloid (0.717 vs. 0.677; Figure 2C), so we treat Geci as an independent-provenance stress test rather than as a claim of uniform state-of-the-art ranking. Leakage-positive controls are not used for model ranking: ADMET-AI DILI and DILIPred-DILIst-FDA are retained only as overlap controls in the released per-compound files because their training targets intersect the TDC/DILIst-FDA label lineage [59, 61]; ADMET-AI DILI rises to 0.927 on TDC, whose labels overlap its training set.

#### Model identity and channel attribution

Because the proxy-DILI features are themselves DILIPredictor-derived, a natural concern is that Oakuloid’s lead could reflect information inherited from DILIPredictor. We therefore attributed the structural lead across the stack’s three channels by re-fitting the logistic meta learner on nested subsets of the *same* out of fold base scores (Table 1). The gradient boosted backbones carry most of the signal: the strongest single backbone (CatBoost) already reaches AUROC 0.807 on strict external TDC, above every evaluated baseline (DILIPred-Preclin 0.752), and stacking all three GBDTs holds external TDC AUROC at 0.805 (Table 8). The benchmark lead is therefore primarily attributable to the gradient boosted descriptor models rather than to the added channels. Adding the four fingerprint random forests is approximately AUROC neutral (+0.001 on TDC, −0.012 on the fully clean TDC subset; they are retained for decision-boundary diversity), and the four predicted proxy-DILI features add a small further increment (+0.005 on TDC, +0.008 on the clean TDC subset).

**Table 1.** Channel-attribution ablation. The logistic meta learner is re-fit on nested subsets of the *same* out of fold base scores (seed-ensemble AUROC): M0, the three GBDT backbones; M1, + the four fingerprint random forests; M2 (Oakuloid), + the four predicted proxy-DILI features. The gradient boosted backbones already exceed every evaluated baseline (CatBoost alone reaches 0.807 on TDC; DILIPred-Preclin 0.752); the fingerprint forests are approximately neutral, and the proxy features add a small increment that persists on the provenance clean doubly disjoint Geci set, indicating the DILIPredictor-derived proxy channel does not account for the headline lead. M2 reproduces the shipped model to within seed-ensemble rounding (TDC 0.811, TDC clean 0.790, and Geci doubly-disjoint 0.590 match the headline values used in this section).

| Stack variant | TDC ext. ($n=401$) | TDC clean ($n=85$) | Geci doubly-disj. ($n=46$) |
| --- | --- | --- | --- |
| M0: three GBDT backbones | 0.805 | 0.793 | 0.581 |
| M1: + four fingerprint RF | 0.806 | 0.781 | 0.579 |
| M2: + four proxy-DILI (Oakuloid) | 0.811 | 0.790 | 0.590 |

Because those proxy features are DILIPredictor-derived and share label provenance with public benchmarks, we verified their contribution on the doubly disjoint, provenance clean Geci set, where they add +0.010 AUROC (0.590 vs. 0.579). This directionally positive gain on the clean-provenance set does not account for the headline lead, so the comparison against DILIPredictor does not depend on the proxy channel. A direct leakage audit of the channel reinforces this interpretation: each cross-endpoint proxy model excludes every DILI evaluation compound from its own training set, with zero deployed InChIKey-14 overlap across TDC-401 and Geci despite substantial raw structural overlap (e.g. 288 shared TDC compounds for the clinical-toxicity endpoint), and the proxy scores correlate only weakly with the DILI label (Pearson *r* ≈ 0.20). The channel is therefore a weak leakage-clean additive signal, not a direct DILI-label surrogate.

### 2.2 Leakage audit and structural generalization limit

The leakage audit asks how much of the ordinary external performance survives after removing increasingly plausible sources of benchmark advantage. It should not be read as a single monotonic curve across unrelated datasets. The ordinary cross-dataset comparison in Figure 2 is therefore separated from the leakage audit in Figure 3: the TDC full, scaffold-disjoint and fully clean subsets test increasingly strict filters within the same TDC mother dataset, whereas the Geci columns test transfer to independent label provenance. Isolating the TDC ladder (Figure 3A), Oakuloid remains ahead from the full strict external set (0.811, *n* = 401) through the scaffold disjoint set (0.755, *n* = 200) and the fully leakage clean TDC subset (0.790, *n*_clean_ = 85). On the fully clean TDC subset, DILIPred-Preclin is 0.655 (standalone margin +0.135); on the paired *n* = 84 subset used for DeLong testing (one compound is dropped from the 85-compound clean subset for lack of a paired comparator score) the difference is +0.156 with *p <* 0.001.

**Fig. 3.**
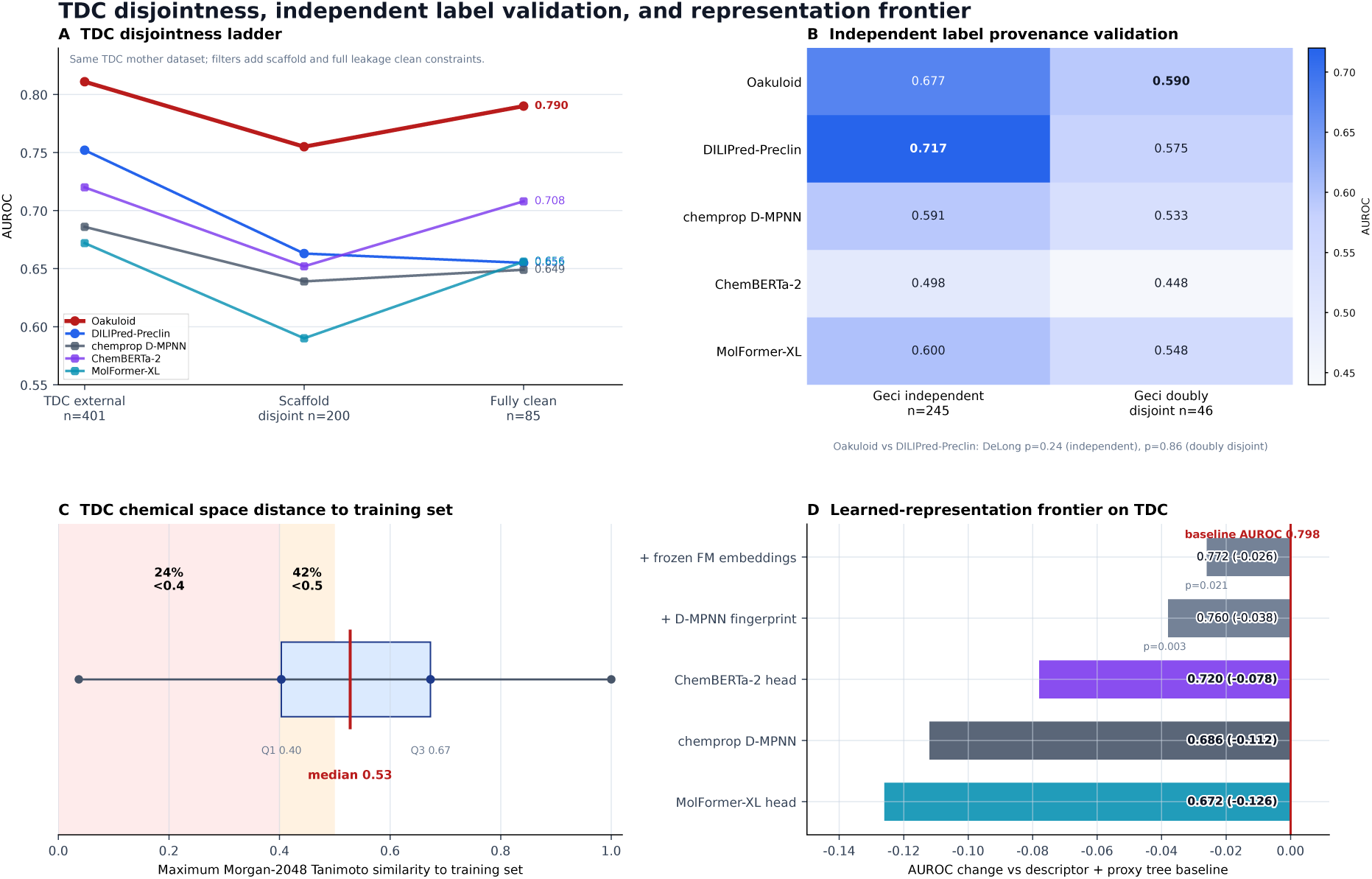
TDC disjointness ladder, independent label validation and learned representation frontier. **(A)** Within TDC AUROC ladder: strict external, scaffold disjoint and fully leakage clean subsets. **(B)** Geci independent label provenance validation, shown separately from the TDC ladder. **(C)** TDC chemical space distance to the Oakuloid training set, summarized by nearest-neighbour Morgan-2048 Tanimoto similarity. **(D)** Controlled learned representation ablations on the TDC external benchmark.

The Geci validation changes the interpretation from a simple ranking to an applicability-boundary analysis. In the larger Geci independent set, the label provenance shifted comparator DILIPred-Preclin edges ahead (0.717 vs. Oakuloid 0.677). On the smallest and hardest set, the Geci doubly disjoint cohort [21] (*n* = 46), we do not test all pairwise contrasts: DILIPred-HumanHep is numerically highest (0.679) and Oakuloid (0.590) is not significantly different from DILIPred-Preclin (0.575; DeLong *p* = 0.86), so this set is too small to support stable ranking and is best interpreted as a conservative lower-bound check rather than a decisive margin for any method.

The same boundary appears within the TDC benchmark. Here, 24% of compounds have maximum Morgan-2048 Tanimoto similarity *<* 0.4 to the training set and the median nearest neighbour similarity is 0.53. Oakuloid’s +0.06 AUROC lead over DILIPred-Preclin is concentrated in the training-similar regime (maximum Tanimoto ≥ 0.4, *n* = 304: 0.846 vs. 0.771, +0.075) and is not observed on the 97 structurally novel compounds with maximum similarity *<* 0.4, where both methods fall to near-chance discrimination (Oakuloid 0.654, DILIPred-Preclin 0.650; fully overlapping bootstrap confidence intervals). The benchmark lead therefore reflects coverage of the training distribution rather than a generalization advantage on structurally novel compounds. This within-TDC result is consistent with the conservative Geci result and clarifies the model’s applicability domain. The accompanying physicochemical comparison is reported in Supplementary Table S9.

The representation frontier analysis gives a similar message. In a controlled TDC ablation, the descriptor-plus-proxy assay tree baseline reaches AUROC 0.798; adding frozen MolFormer/ChemBERTa embeddings or a supervised D-MPNN fingerprint does not improve it (AUROC 0.772, Δ = −0.026, *p* = 0.021; and to 0.760, Δ = −0.039, *p* = 0.003), and the end-to-end learned heads land a further ∼ 0.08–0.13 lower (Supplementary Table S2). Thus, in this benchmark setting, richer learned representations do not improve over the descriptor-plus-proxy tree. The full stack used as Oakuloid reaches 0.811 on the same full TDC benchmark; the 0.798 value is the matched ablation baseline used only for representation comparisons. Added directly as base learners to the full Oakuloid stack, the same frozen embeddings shift TDC AUROC negligibly (MolFormer +0.003, ChemBERTa +0.000, both +0.004; within the five-seed noise band), so this conclusion holds for the deployed stack and is not an artifact of the matched-base comparison.

### 2.3 Internal structure-model baseline, BIO wet lab signal and structure/biology fusion

This section starts with the structure-only reference point on the internal DXTox/DILIrank wet lab cohort, then evaluates the iBAC BIO signal, rank-mean fusion and net-benefit analysis. The wet lab analysis uses the internal DILIrank cohort, so the structure-only result on this same cohort is the relevant comparator. On the *N* = 94 binary subset, Oakuloid reaches AUROC **0**.**850** (AUPRC 0.941, MCC 0.611; five seed mean 0.844 ± 0.010), while DILIPred-Preclin reaches 0.832 and is statistically indistinguishable on this small cohort. The two chemical language model baselines are near random on the same compounds (MolFormer-XL AUROC 0.483, MCC 0.115; ChemBERTa-2 0.497, MCC 0.196). At the internal Youden threshold (0.604), Oakuloid favours sensitivity, flagging 37*/*38 vMost-DILI and 29*/*33 vLess-DILI compounds at 15*/*23 vNo-DILI specificity, whereas DILIPred-Preclin favours specificity (sensitivity/specificity 0.817*/*0.783).

An orthogonal, directly measured signal can refine this operating point beyond what structure alone provides. We therefore measured the iBAC 3D-PHH IC_50_/C_max_ ratio (BIO) for the 122-compound DILIrank wet lab cohort and evaluated the *N* = 94 binary subset. BIO alone reaches AUROC 0.758, with high specificity (21*/*23 = 0.91) but limited coverage of less severe DILI (8*/*33 = 0.24 vLess-DILI sensitivity). The Youden threshold for the BIO score corresponds to IC_50_/C_max_ ≤ 36.77; the single compound exactly on the threshold is assigned to the positive side; under this convention, BIO captures 31*/*38 = 0.82 vMost-DILI compounds. A single inhibition-rate measurement at 30 × *C*_max_ provides a moderate surrogate for the full concentration-response signal among covered compounds (AUROC 0.742, Spearman *ρ*_3-level_ = 0.469; Figure 4).

**Fig. 4.**
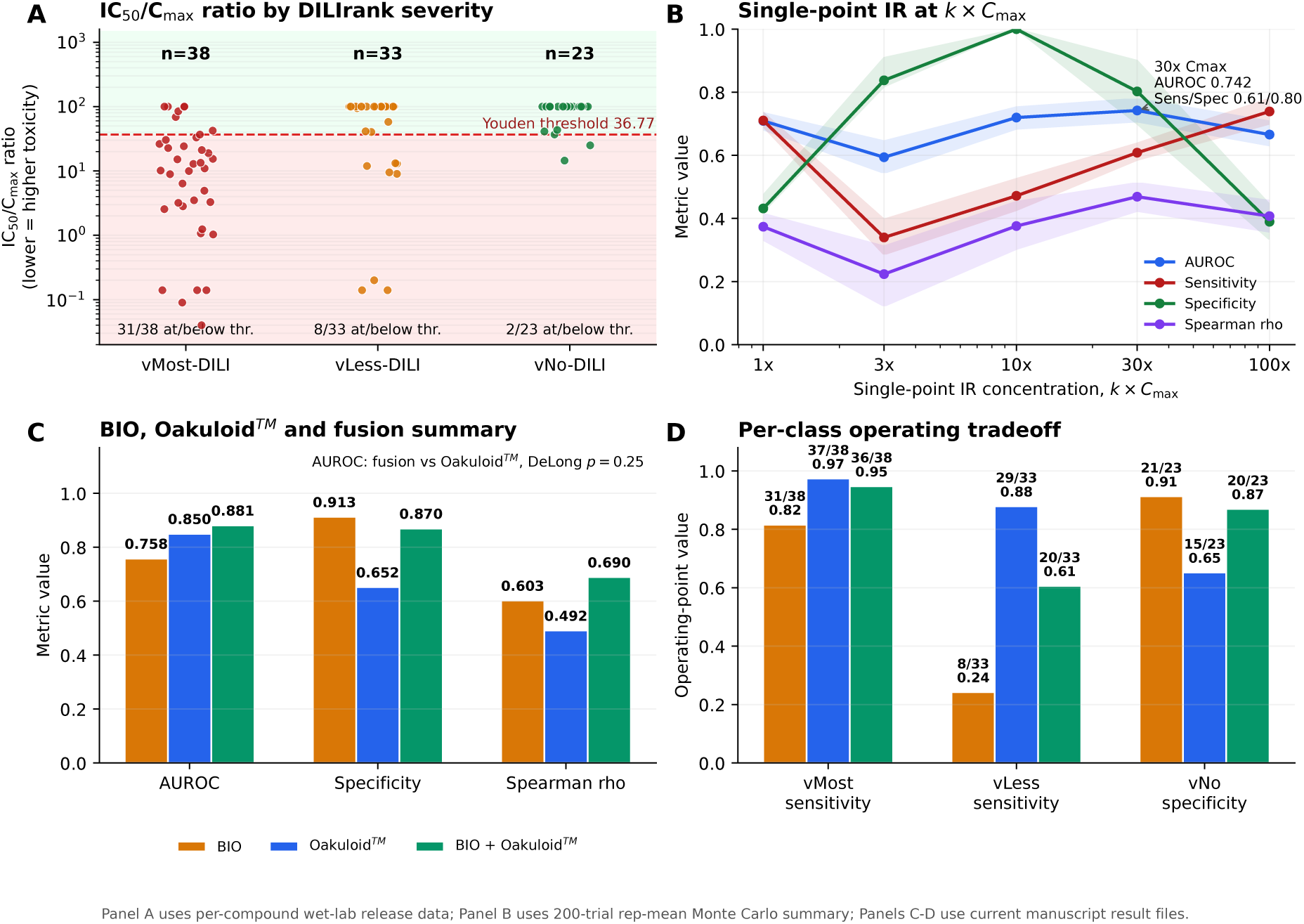
Wet lab BIO signal and structure/biology fusion. **(A)** IC_50_/C_max_ values by DILIrank class, with the Youden threshold at 36.77. **(B)** Single-point inhibition-rate performance across *k×C*max concentrations. **(C)** AUROC, vNo specificity and three-level Spearman *ρ* for BIO, Oakuloid and rank mean fusion on the *N* = 94 binary cohort. **(D)** Per-class operating point summary, showing that fusion mainly trades vLess-DILI sensitivity for improved vNo-DILI specificity.

The wet lab and structural channels are complementary but not interchangeable. A structure only regressor cannot reconstruct the measured IC_50_/C_max_ ratio (*R*^2^ = −0.020, Spearman 0.31); replacing the measured ratio with a structure-predicted ratio costs 0.088 AUROC. Rank-mean fusion of Oakuloid and measured BIO raises internal cohort AUROC from 0.850 to 0.881, significantly above BIO alone (*p* ≈ 0.001) but not significantly above Oakuloid alone (DeLong *p* = 0.25). The more robust interpretation is therefore not an AUROC gain over the structural model, but a change in operating mode: specificity rises from 15*/*23 = 0.65 to 20*/*23 = 0.87, while vMost-DILI sensitivity remains high (36*/*38 = 0.95) and vLess-DILI sensitivity falls from 29*/*33 = 0.88 to 20*/*33 = 0.61.

A decision-curve (net-benefit) analysis [16] makes this operating-mode reading precise (Figure 5). Across a pre-specified grid of threshold probabilities, structure-plus-BIO fusion does *not* improve on the structural model in net-benefit terms on the internal cohort; the structural score has higher net benefit across most of the grid, with fusion higher only in a narrow high-threshold window. We therefore do not claim a fusion benefit beyond the operating-point shift. On strict-external TDC, by contrast, Oakuloid’s net benefit equals or exceeds DILIPred-Preclin’s across 82% of the grid and remains net-beneficial to a higher threshold (*p*_t_ ≈ 0.77 vs. 0.72). In both cohorts the models’ incremental value over treat-all/treat-none is realized at high decision thresholds, corresponding to confirm-before-escalate decisions, rather than as a discrimination gain.

**Fig. 5.**
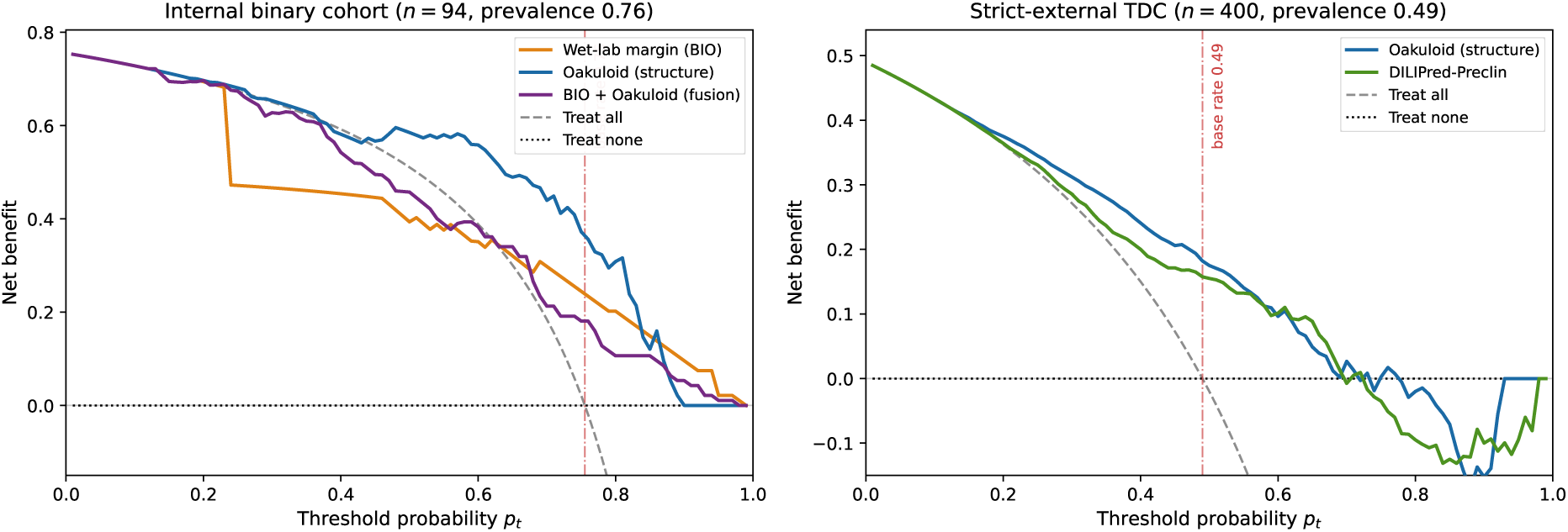
Decision-curve (net-benefit) analysis. Net benefit versus threshold probability on a pre-specified grid (flag if predicted risk *≥ pt*), with treat-all and treat-none references. **(A)** Internal DILIrank cohort (*N* = 94): structure-plus-BIO fusion does not exceed the structural model in net benefit across most thresholds; the structural score is generally higher, with fusion ahead only in a narrow high-threshold window. **(B)** Strict-external TDC (*n* = 400): Oakuloid net benefit equals or exceeds DILIPred-Preclin across the majority of the grid and remains beneficial to a higher threshold. In both panels every model curve stays net-positive in the high-threshold regime where “treat-all” has turned negative, locating the models’ value in confirm-before-escalate decisions rather than in a discrimination (AUROC) gain.

The fusion rule itself is a single equation. For a structural score *s* ∈ [0, 1] and wet lab ratio *r >* 0, the deployed score is

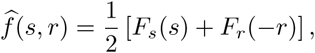

where *F*_s_ and *F*_r_ are empirical CDFs of structural scores and negated wet lab ratios. Cohort-rank fusion gives AUROC 0.881; the fixed CDF version for single-compound inference gives 0.879, making it practically equivalent for deployment. Per compound inspection also clarifies the tradeoff: Clozapine is correctly reclassified by the wet lab channel, Simvastatin remains correctly called by structure and fusion, whereas Acetaminophen, Labetalol and Alendronate illustrate cases where a high IC_50_/C_max_ can shift a true DILI compound below the fused threshold. These cases are released in Supplementary Data 4.

### 2.4 Clinical stage DILI failures

This panel is a sensitivity stress test on known positives, not a benchmark or a specificity claim. We evaluated 13 drugs that failed clinical development or were withdrawn because of hepatotoxicity. At the external set Youden threshold (0.733), Oakuloid flags all 13*/*13 cases (Table 2); DILIPred-Preclin also flags all 13 at its native 0.5 cutoff. Thus, the clinical panel supports sensitivity, but not method-specific superiority. Three observations make the result more informative than the aggregate count alone. First, the recall is consistent with the model’s structural evidence: 11*/*13 carry at least one of the reactive metabolite or mitochondrial structural alerts the model relies on (Section 2.6); the two that carry none (troglitazone, whose quinone-methide is not captured by coarse MACCS keys, and evobrutinib) are still scored above threshold, indicating that the model is not a simple alert-matching rule. Second, the conformal layer (Section 2.5) distinguishes *how* confidently each is flagged: the top seven are confident singletons ({DILI}, confidence 0.90–0.95), whereas the lower six, including the closest call, evobrutinib at 0.748, fall in ambiguous {both} sets, a direct indication that not all recovered positives are equally certain. Third, although both methods reach 13*/*13, Oakuloid scores higher than DILIPred-Preclin on 11*/*13 and by +0.16 on average on the three animal-negative idiosyncratic toxicants (fialuridine, troglitazone, firazorexton), where structure based recall is particularly relevant. The 13*/*13 result is not threshold dependent within the evaluated range; it holds for any operating threshold up to 0.748. We nevertheless report it as a sensitivity demonstration on known positives, not a specificity or superiority claim.

**Table 2.**
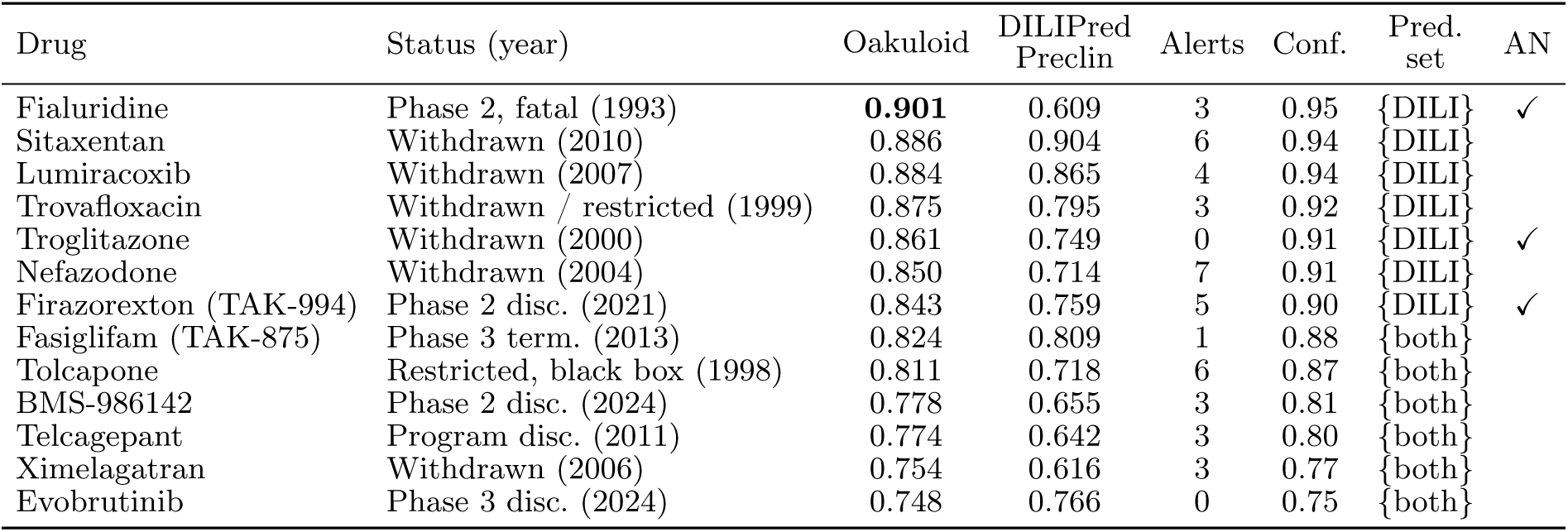
Clinical stage DILI failures and withdrawals. Predicted DILI probabilities; Oakuloid is flagged at the external set Youden threshold 0.733 and DILIPred-Preclin at its native 0.5 cutoff (both reach 13*/*13). “Alerts” counts how many of the model’s enriched reactive metabolite/mitochondrial MACCS structural keys (Figure 7B) the compound carries. “Conf.” and “Set” are the split conformal point confidence and prediction set at *α* = 0.10 ({DILI} = confident DILI singleton; {both} = ambiguous, i.e. non-DILI cannot be excluded); “AN” marks the three strict animal-negative cases. Bold marks the highest Oakuloid score. Scores are from nc_clinical_predictions.csv; per drug detail in Supplementary Table S8.

### 2.5 Calibration, uncertainty and selective prediction

Beyond discrimination, deployment requires a signal for when predictions should be deferred. On the internal cohort, Oakuloid’s absolute discrimination is robust to near-duplicate removal: dropping close internal neighbours changes AUROC only from 0.850 to 0.843 (Tanimoto ≥ 0.85, *n* = 83) and to 0.840 (≥ 0.70, *n* = 70), so the internal result is not driven by near duplicate chemistry (the comparative lead’s similarity dependence is treated separately in Section 2.2). Probability calibration is adequate for ranking but not for unaudited absolute-risk use [11, 22]: the Brier/ECE of 0.136*/*0.150 on DILIrank (*N* = 94), 0.210*/*0.176 on TDC (*n* = 401) and 0.226*/*0.139 on the out-of-distribution Geci set (*n* = 245) reflect mild overprediction relative to the balanced class rate. Post-hoc recalibration does not help here: a Platt fit on the groupwise out-of-fold predictions is a near-identity map. The meta learner is itself logistic, so its out-of-fold scores are already sigmoid-calibrated on the training distribution; Platt scaling leaves Brier/ECE essentially unchanged, while isotonic regression overfits the out-of-fold scores and degrades both calibration and ranking. We therefore report Oakuloid probabilities uncalibrated and attribute the residual external miscalibration to the train-versus-cohort prevalence shift, which a single global scalar cannot absorb. Reliability diagrams for both cohorts are shown in Supplementary Figure S1.

Operating-point metrics, by contrast, must be reported without tuning the threshold on the evaluation cohort. To pre-empt this optimistic-threshold concern we fixed a single decision threshold on the 4,681-compound training out-of-fold predictions (*τ* = 0.694, maximizing training MCC; never exposed to any external label) and applied it unchanged to every external cohort. At this leakage-clean fixed operating point Oakuloid attains MCC 0.480*/*0.540*/*0.364 and balanced accuracy 0.737*/*0.808*/*0.669 on TDC-401 / internal DILIrank / Geci-245, a modest 0.019–0.062 MCC reduction from the per-cohort own-Youden optima (0.526*/*0.602*/*0.383). This expected reduction from avoiding test set threshold tuning confirms that the ranking performance is not an artefact of an over-fit operating point. A sensitivity-weighted clinical threshold (*τ* = 0.351, 2:1 cost) raises sensitivity to ≥ 0.98 on all three cohorts at the expected specificity cost.

To turn this into a deployable abstention signal, we wrap Oakuloid in a split conformal predictor with class conditional (Mondrian) calibration, which gives finite-sample coverage under exchangeability [4, 65]. Empirical coverage matches the nominal target at every level and holds for both classes (Table 3); lower nominal coverage targets shrink the prediction set toward a single label. The same confidence enables a selective prediction mode: ranking compounds by conformal confidence and abstaining on the least confident fraction raises AUROC from 0.813 at full retention to 0.845 at 70% (Figure 6), defining a risk-tiered workflow in which low confidence out-of-domain compounds are routed to the orthogonal iBAC assay. As an out-of-distribution robustness probe, recalibrating the conformal layer on training scores and applying it to the Geci set shows that coverage deteriorates when exchangeability is violated (negative-class coverage falls to 0.64/0.54 at *α* = 0.1*/*0.2). The abstention layer therefore provides a practical route for sending low-confidence out-of-domain compounds to the assay.

**Fig. 6.**
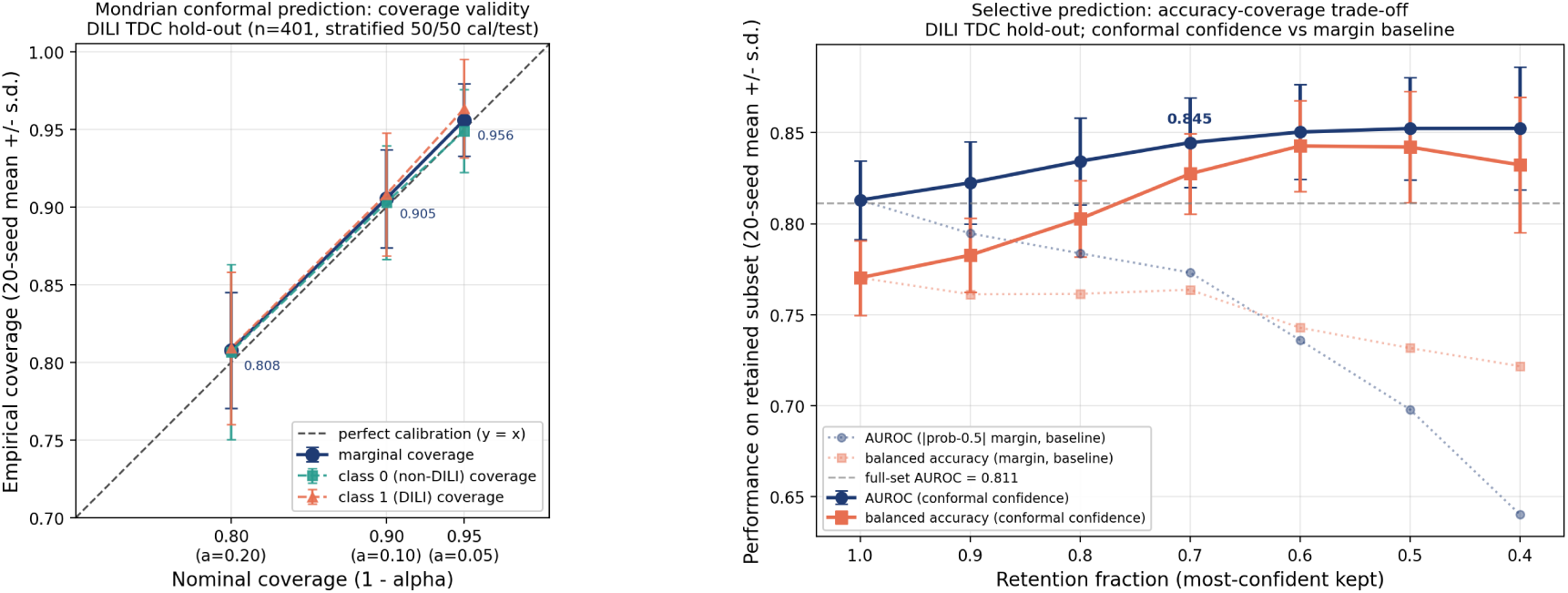
Calibrated uncertainty. **(A)** Split conformal empirical coverage matches the nominal target at every level, marginally and class conditionally. **(B)** Selective prediction by conformal confidence: AUROC rises monotonically as the least confident compounds are abstained on and routed to the orthogonal wet lab assay, from 0.813 at full retention to 0.845 at 70%, staying above the probability margin baseline.

**Table 3.** Conformal coverage and selective prediction efficiency. Split conformal prediction with class conditional (Mondrian) calibration on the held out TDC predictions (20 random calibration/test splits). For each target error rate *α*, empirical marginal coverage matches the nominal 1 *− α* and is also maintained class conditionally for both DILI positive (class 1) and DILI negative (class 0) compounds; lower nominal coverage targets yield smaller prediction sets (mean set size toward 1, higher singleton rate). Selective prediction by conformal confidence raises AUROC from 0.813 at full retention to 0.845 at 70% retention (Supplementary Table S5; Figure 6).

| Nominal $1 - \alpha$ | Marginal cov. | Class 0 cov. | Class 1 cov. | Mean set size | Singleton rate |
| --- | --- | --- | --- | --- | --- |
| 0.95 | 0.956 | 0.949 | 0.963 | 1.69 | 0.31 |
| 0.90 | 0.905 | 0.903 | 0.908 | 1.45 | 0.55 |
| 0.80 | 0.808 | 0.807 | 0.809 | 1.14 | 0.86 |

### 2.6 Mechanistic interpretability

SHAP [42] and MACCS structural-alert enrichment are used as transparency checks rather than as causal mechanism discovery (Figure 7). After the leakage clean retrain the twenty highest-SHAP features are all interpretable molecular descriptors, including BCUT charge/ionisation eigenvalues, partial charge surface areas and autocorrelation descriptors, with no training source identity (src_) bit in the ranking (Figure 7A); in leakage-prone configurations a source identity bit instead dominated the attribution. The enrichment analysis recovers recognised reactive metabolite and mitochondrial chemotypes among DILI positive compounds (nitroaromatics, hydrazine/azo *N* –*N*, aromatic halides and nitriles; odds ratios ≈ 2–3, all FDR *q <* 0.05), while flagging the organosilicon key as a training set specific artifact rather than a mechanistic alert (Figure 7B). A compact subset of the enriched keys is shown in Table 4 so that the main mechanistic and artifact calls can be evaluated directly in the main text. The complete enrichment table and the top twenty clean model SHAP table are provided in Supplementary Tables S6–S7; the full 862 feature SHAP and source identity audit is released as Supplementary Data 2.

**Fig. 7.**
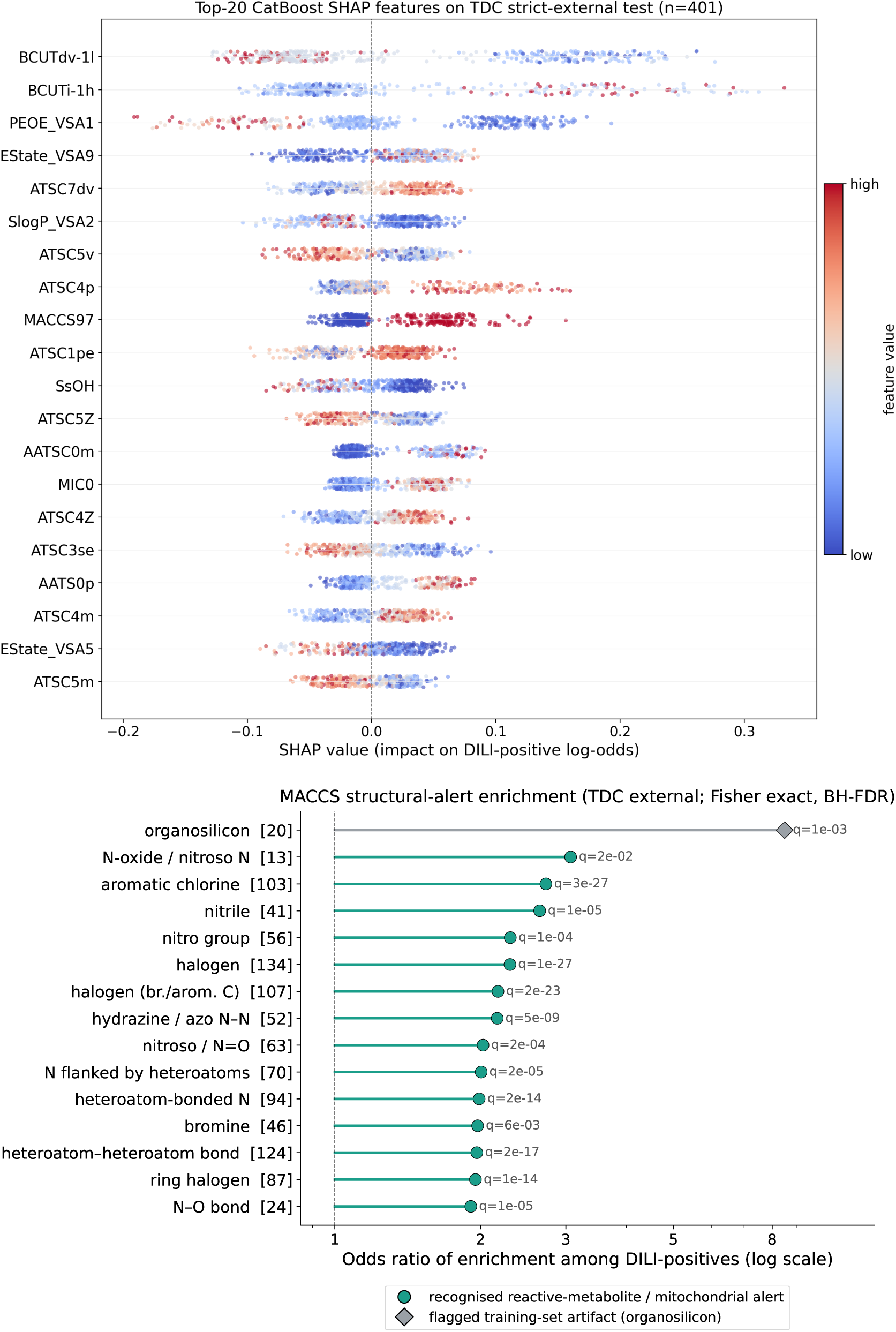
Mechanistic interpretability of the leakage clean model. **(A)** SHAP attribution of the leakage clean CatBoost backbone on the strict external TDC set (*n* = 401): all twenty highest-impact features are interpretable molecular descriptors and none is a training source identity (src_) bit, in contrast to the leakage prone configuration in which such a bit dominated the same ranking. **(B)** Odds-ratio forest plot of MACCS structural key enrichment among DILI positive compounds (Fisher’s exact test, Benjamini–Hochberg FDR *q <* 0.05): recognised reactive metabolite and mitochondrial chemotypes are significantly overrepresented, whereas the organosilicon key (rightmost) is annotated as a training set composition artifact and excluded from mechanistic interpretation.

**Table 4.** Representative enriched structural alerts. Compact main text summary of the MACCS structural keys underlying Figure 7B. Bit 20 is retained as an audit flag because it is statistically enriched but is excluded from mechanistic interpretation as a training set composition artifact. The complete 15 key enrichment table is in Supplementary Table S6.

| MACCS bit | Alert class | OR | FDR $q$ | $n$ | Interpretation |
| --- | --- | --- | --- | --- | --- |
| 20 | Organosilicon | 8.49 | $1.1 \times 10^{-3}$ | 22 | Training set composition artifact |
| 13 | $N$-oxide / nitroso type $N$ | 3.07 | $1.5 \times 10^{-2}$ | 33 | Reactive nitrogen alert |
| 103 | Chlorine / aromatic halide | 2.73 | $2.5 \times 10^{-27}$ | 766 | Reactive metabolite associated halide |
| 41 | Nitrile | 2.65 | $1.3 \times 10^{-5}$ | 126 | Cyanide release / reactive chemistry |
| 56 | Nitro group | 2.30 | $9.8 \times 10^{-5}$ | 132 | Nitroaromatic and mitochondrial risk |
| 52 | Hydrazine / azo $N-N$ | 2.16 | $5.2 \times 10^{-9}$ | 339 | Classic reactive metabolite alert |

**Table 5.** Deployment decision guide. Intended use of the four Oakuloid operating modes, with internal-cohort (*N* = 94) operating points reported in the text. The conformal-abstention row reports TDC selective prediction.

| Mode | Threshold / source | Intended use | Operating point and caveat (internal $N = 94$) |
| --- | --- | --- | --- |
| Structure-only screen | Youden (internal 0.604, external 0.733) | Broad first-pass triage of any compound with a SMILES | High sensitivity (vMost 37/38, vLess 29/33) at vNo specificity 15/23; flags additional non-DILI compounds |
| Wet-lab BIO alone | $IC_{50}/C_{max} \leq 36.77$ | Orthogonal cytotoxicity readout where assayed | High specificity (21/23) but low coverage of mild DILI (vLess 8/33) |
| Rank-mean fusion | rank-mean(structure, -ratio) | Higher-specificity confirmation for compounds with a measured ratio | Specificity 20/23, vMost 36/38; trades vLess sensitivity (20/33); not a significant AUROC gain over structure ($p = 0.25$) |
| Conformal abstention | lowest split-conformal confidence | Triage: route low-confidence compounds to the iBAC assay | AUROC 0.813 → 0.845 at 70% retention |

## 3 Discussion

Oakuloid addresses two recurring problems in computational DILI prediction: standard external benchmarks often retain compound or label provenance overlap, and structure only models cannot observe exposure- and biology-dependent injury. Evaluated under a leakage clean protocol, the four proxy stack has the highest AUROC on the TDC benchmark and the within TDC disjointness ladder. The independent Geci validation gives a more conservative provenance picture: on the smallest doubly disjoint set, no stable ranking is established (Oakuloid is not significantly separated from DILIPred-Preclin, *p* = 0.86; DILIPred-HumanHep is numerically highest), and on the larger Geci independent set a preclinical comparator is numerically higher. A measured wet lab signal then adds orthogonal specificity to the structural screen. Recent DILI work has advanced model classes and assay fidelity [27, 34, 43, 46, 78], but has less often combined released per-compound predictions, a multi-axis leakage audit with independent label-provenance validation, and a paired structure/wet-lab screen-and-confirm workflow. Oakuloid contributes this combination rather than a single new model class or assay.

The wet lab result is therefore best interpreted as evidence of orthogonality, not as evidence that fusion is uniformly superior to the structural model. On AUROC, BIO + Oakuloid is not significantly higher than Oakuloid alone (0.850 → 0.881, DeLong *p* = 0.25). Its practical value is an operating point shift: the structure model is a high sensitivity screen, whereas fusion is a higher specificity confirmation mode that can rule out non-DILI compounds overflagged by the structural score. That benefit comes with an explicit tradeoff: less severe or type-B DILI compounds with weak 72-hour cytotoxicity can fall below the fused threshold. The recommended use is therefore to score broadly with the structure model and reserve the iBAC assay for borderline or high-value compounds where an orthogonal biological measurement can change a decision. This orthogonality is independently corroborated beyond our assay: a recent transcriptomics model trained on primary human hepatocyte expression (DILImap) [8] attains high specificity on a blind compound set from measured biology rather than structure, supporting measured cellular signal, whether the iBAC IC_50_/C_max_ ratio or expression, as a complementary axis to structural prediction and a natural future fusion partner.

Several limitations remain. The 4,681-compound training corpus is still small by modern representation-learning standards and inherits animal-model and FDA/LTKB label provenance biases from public DILI resources. The internal wet lab cohort is the assay-development cohort for the iBAC platform, so fusion performance should be treated as single-site evidence rather than as a held out wet lab validation. The internal cohort was also filtered from a 143-compound master table by procurement and solubility constraints and skews toward clinically prominent, well-characterized drugs, a selection bias that may inflate apparent internal performance. The model is tuned for sensitivity and is not perfectly calibrated on the balanced external set; threshold dependent metrics are reported as operating points, while AUROC and ordinal Spearman are the main comparative metrics. Finally, the representation frontier analysis uses frozen Transformer embeddings and controlled ablations; it does not exclude future gains from substantially larger labelled DILI datasets or task-specific pretraining. Relatedly, the benchmark lead is similarity dependent: as the within-TDC similarity stratification shows (Section 2.2), Oakuloid’s advantage over the baselines is concentrated on training-similar chemistry and narrows on the most structurally novel compounds. The headline AUROCs should therefore be read as benchmark performance, whereas generalization to structurally novel chemistry remains constrained for all evaluated structure-only methods.

The most direct next step is to expand the iBAC IC_50_/C_max_ dataset beyond the present 122-compound DILIrank panel and validate the fixed CDF fusion rule on a prospectively held out wet lab cohort. The more difficult step is to add signal classes that a single 72-hour cytotoxicity endpoint cannot capture, such as immune-cell co-culture, transcriptomics or human genetic susceptibility. Those measurements are most likely to improve the type-B and chronic injury cases that remain difficult for both structure and cytotoxicity.

## 4 Conclusions

Oakuloid provides an open, leakage audited DILI workflow that pairs a four proxy structural ensemble with an optional iBAC IC_50_/C_max_ measurement: a high sensitivity structural screen that has the highest AUROC among the out-of-distribution-valid baselines on the TDC benchmark and within TDC disjointness ladder, and a rank mean fusion rule that converts an orthogonal wet lab measurement into a higher specificity confirmation mode. Throughout, the strict external TDC result is the primary generalization estimate, while the internal DILIrank cohort is an in-domain reference that shares FDA/LTKB label provenance with training. Released with the model are the provenance-disjoint benchmark, the iBAC wet lab panel, and a model card, providing a shared leakage audited reference for deployment studies and future measured signal classes.

## 5 Materials and methods

### 5.1 Datasets

#### 5.1.1 Internal DILIrank cohort (n = 122)

We assembled 122 marketed and clinical stage drugs spanning small-molecule diversity (MW 100–800 Da, logP −3 to +8). The cohort was constructed from a master 143-compound C_max_ table prepared in our internal program. We retained the 122 entries that satisfied four predefined inclusion rules: (1) availability of an unambiguous PubChem SMILES (RDKit-parseable); (2) commercial availability of a ≥ 95%-purity stock at the time of assay scheduling; (3) solubility in ≤ 0.5% DMSO at the highest planned dose (≥ 30×C_max_); and (4) presence of an FDA DILIrank vDILI-Concern annotation [13]. The Ambiguous and unlabeled categories are recorded but excluded from all binary and ordinal evaluations. Labels were merged by international nonproprietary name after salt-counter-ion normalization.

The 21 compounds dropped from 143 to 122 failed rule (2) or (3) because of procurement or solubility constraints. This filtering was performed before any predictive model was trained or evaluated and is therefore prospective relative to the benchmark. Compounds were not prescreened on the DILIrank label before inclusion. For each retained compound we recorded (a) an in vitro IC_50_ measured on the iBAC 3D-PHH platform (Section 5.2), (b) the curated total plasma C_max_, and (c) the DILIrank vDILI-Concern label. The source pool was assembled over time and is enriched for clinically prominent drugs with abundant pharmacokinetic literature; this selection bias is discussed in the Limitations.

#### 5.1.2 Disclosure of overlap with our 2021 iBAC validation set

The present 122-compound cohort is essentially the same set of clinically tested drugs used to validate the iBAC platform in our 2021 paper [73]; the iBAC platform was first published using this cohort. The DILIrank labels and the IC_50_/C_max_ values used here are not newly held out; they were available to the assay-development team at the time of platform validation. This creates a potential optimism for any platform-vs-DILIrank performance comparison, and we explicitly do *not* claim that the iBAC platform was developed blind to DILIrank. However, the structure only ML side of the fusion (Oakuloid) was trained on a 4, 681-compound compendium deduplicated by InChIKey-14 against the internal cohort and the TDC subset (see Section 5.9), and the strict external TDC validation (Section 2.1) is on a wholly independent benchmark with overlap removed. We caution that the gold standard component of this training set (DILIst/DILIrank) shares FDA/LTKB label provenance with the internal cohort and with TDC, so InChIKey-14 deduplication removes overlapping *compounds*but not the shared label *provenance*; standard benchmark gains are reported as such, and the doubly disjoint novel chemistry result (AUROC ≈ 0.59, Section 2.2) is the provenance-clean lower bound. The fusion AUROC of 0.881 should therefore be interpreted as a development-cohort estimate under our assay system, not as a held out generalization estimate.

The 4-class DILIrank label distribution and per class size of the three evaluation protocols are summarized in Table 6. Ambiguous-DILI-concern (*n* = 12) and unlabeled compounds (*n* = 16) are excluded from all binary and ordinal evaluations.

**Table 6:** Internal DILIrank cohort: 4-class label distribution and the three evaluation protocols. Numbers in parentheses are class sizes (positive / negative).

| DILIRank class | $n$ | Used in protocol |
| --- | --- | --- |
| vMost-DILI-concern | 38 | Binary (pos), Most-vs-No (pos), 3-level ($y = 2$) |
| vLess-DILI-concern | 33 | Binary (pos), 3-level ($y = 1$) |
| vNo-DILI-concern | 23 | Binary (neg), Most-vs-No (neg), 3-level ($y = 0$) |
| Ambiguous-DILI-concern | 12 | excluded |
| unlabeled | 16 | excluded |
| **Total** | **122** | NA |

We evaluate using three orthogonal protocols, in increasing order of stringency:

1. **Binary** (primary): {vMost, vLess} as positive vs. vNo as negative, **N**= **94** (71 pos / 23 neg).
2. **Most-vs-No** (strictest): vMost as positive vs. vNo as negative, *N* = 61 (38 pos / 23 neg). Used for severity-stratified sanity checks.
3. **3-level ordinal** (Spearman *ρ*): vMost = 2, vLess = 1, vNo = 0, *N* = 94. Tests whether the predicted score correlates monotonically with severity rather than only separating two classes.

#### 5.1.3 Concentration response cohort (n = 132)

For the 122-cohort compounds plus additional reference compounds (n = 132 total in the released raw-readout file), concentration-response cellular inhibition rates (IR) were measured following the previously published iBAC protocol [73] (also Section 5.2). The full per-well readout file (1,909 rows, 132 compounds) is released as Supplementary Data 5.

#### 5.1.4 External TDC DILI subset (n = 401)

We benchmarked on the Therapeutics Data Commons (TDC) DILI task (Huang et al. 2021) [28], *n* = 475. To avoid evaluation leakage, we removed any compound matching an InChIKey-14 in either (a) the Oakuloid training set (33/475 removed, 6.9%) or (b) our internal 122-cohort (41/475 removed, 8.6%); the two InChIKey-14 removal sets are disjoint, so the combined removal is 33 + 41 = 74 compounds and 401 remain. The strict external subset contains **401 compounds** (196 pos / 205 neg), all unseen by the Oakuloid training set and the internal cohort.

#### 5.1.5 Training data for Oakuloid

We use the LivTox proxy-DILI compendium [55], a curated aggregation of nine separate hepatotoxicity datasets, each carrying a numeric *training source identifier* (column Source_rank in the upstream release; values 1–16). We retain only the five sources whose labels are direct DILI-incidence calls (binary, 1 = drug observed to cause clinical or preclinical hepatotoxicity, 0 = otherwise): sources 3 and 7 from Mulliner et al. human and preclinical hepatotoxicity sets (FDA / regulatory drug label review), source 5 from Liu et al. via ToxRefDB [68] (retrospective rodent toxicity study aggregation), source 6 from Ambe et al. via the Japanese Hazard Evaluation Support System (HESS), and source 8 from Shuaibing et al. (large-scale published DILI list); per source counts are summarised in Table 7, and the canonical upstream citation for each source is collected in the LivTox release [55]. The remaining four sources are excluded because they carry *mechanistic-endpoint* labels rather than direct DILI labels: source 14 marks bile salt export pump (BSEP) inhibitors [62], source 15 marks mitochondrial-toxicity hits, and source 16 marks reactive metabolite formers [55]; although each mechanism is a well-established precursor of clinical DILI, the underlying labels answer different prediction questions (e.g. “is this molecule a BSEP inhibitor at ≥ 25 *µ*M IC_50_?”) and mixing them into the DILI training signal would teach the model to predict mechanism rather than DILI incidence. (Mechanistic endpoints are used as auxiliary features in the upstream DILIPredictor design [55], but kept out of our binary-DILI label stream.) Filtering to the five DILI-labelled sources yields 5,646 compounds with valid SMILES (vs. 2,187 for the identifier-=7-only Preclinical subset used by DILIPredictor); the per source breakdown is given in Table 7. To this proxy-DILI base we add the DILIst/DILIrank gold standard clinical reference set, then deduplicate the whole training pool by InChIKey-14 against the internal 122-compound cohort and the TDC subset (the Geci provenance check of Section 5.1.6 is filtered separately), yielding a final **4,681**-compound training set that shares no compound with the internal cohort or the strict-external TDC benchmark. We caution that this gold standard augmentation shares FDA/LTKB label provenance with the internal cohort and with TDC, so InChIKey-14 deduplication removes overlapping compounds but not the shared label provenance; accordingly we report standard benchmark gains as such, and treat the doubly disjoint novel chemistry result (AUROC ≈ 0.59) as the provenance-clean generalization estimate. Per-source positive class rate varies substantially across the five proxy-DILI sources (0.34–0.75), motivating the source identity feature channel introduced in Section 5.3.

**Table 7:**
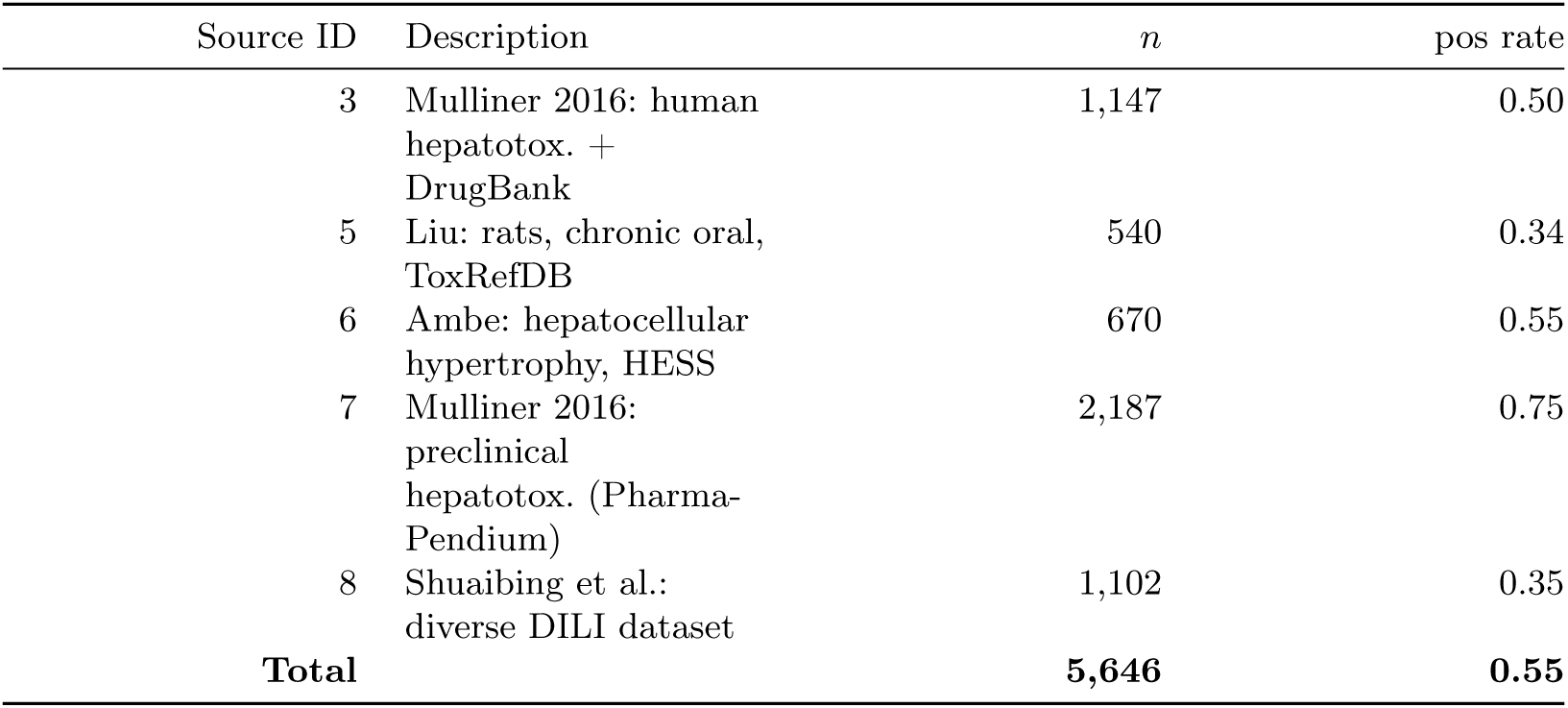
Oakuloid proxy-DILI training source breakdown (5,646 compounds before gold standard augmentation and InChIKey-14 deduplication; the final training set is 4,681 compounds). Sources are taken from the LivTox proxy-DILI dataset; full citations in Seal et al. 2024 [55].

**Table 8.**
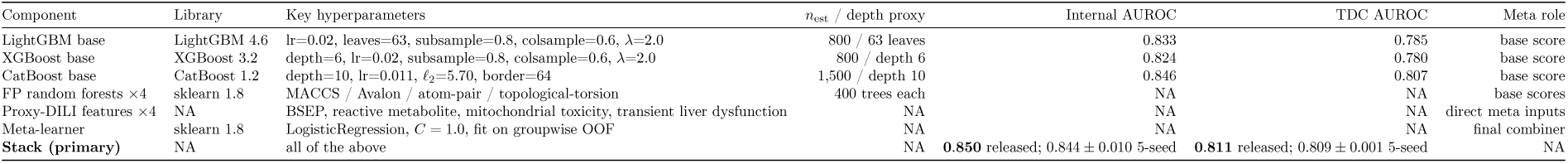
Oakuloid model identity. The primary model stacks three GBDT backbones with four fingerprint random forests and four LivTox proxy-DILI features via a logistic regression meta learner. CatBoost hyperparameters were selected by groupwise (InChIKey-14) cross validated Optuna search on the training set only; the remaining base learners use fixed class balanced configurations. Internal/external AUROC columns are held out seed-ensemble evaluation results, not tuning objectives; the separate leakage-clean CatBoost used only for interpretability (Figure 7) scores 0.767 on TDC and is not the deployed full-feature backbone tabulated here.

#### 5.1.6 Geci independent validation set (n = 245; doubly disjoint n = 46)

For an independent label-provenance check we used the hepatotoxicity dataset of Geci et al. [21], which derives a binary DILI label from exposure-adjusted points of departure rather than from the FDA/LTKB lineage underlying DILIrank, DILIst and TDC. Retaining compounds with a parseable SMILES and a defined label yields **245** evaluable compounds (146 positive / 99 negative; the *Geci independent* set), scored by every method with no model retrained. Because this set shares no FDA/LTKB label provenance with our training corpus, it probes transfer to an independently labelled distribution. We further define a *doubly disjoint* subset of **46** compounds (30 positive / 16 negative) whose InChIKey-14 is absent from the union of all evaluated models’ training corpora. Oakuloid’s training set is a subset of the DILIPredictor corpus, making the DILIPredictor corpus the binding constraint for this filter. This subset is simultaneously novel chemistry and independent label provenance, and provides the conservative provenance-clean generalization estimate (AUROC ≈ 0.59) reported in Section 2.2.

### 5.2 In vitro DILI assay (BIO IC_50_/C_max_ pipeline)

The IC_50_ values used throughout this work were generated on our previously published *integrated biomimetic array chip* (iBAC) platform. Full cell-sourcing, 3D primary human hepatocyte (3D-PHH) culture, hepatic-functionality QC, and 12-day stability characterization details are described in the original iBAC platform report [73].

#### 5.2.1 Cohort assay execution

The 132-compound concentration-response cohort (the 122 DILIrank drugs plus additional reference compounds) was assayed using the iBAC dose-response protocol without modification; dosing scheme, viability readout, and IC_50_-fitting procedure follow the established iBAC platform exactly as previously published [73]. The full per well readout (1,909 rows across 132 compounds) is released as Supplementary Data 5.

#### 5.2.2 C_max_ table

Therapeutic C_max_ values for these 122 compounds were curated from drug label PK sections, FDA review documents, and the literature; the underlying master table from which the cohort was selected contains 143 compounds in total (the 21 unselected compounds failed the inclusion rules above). The IC_50_/C_max_ ratio reported here uses total (not unbound) C_max_ to match the convention used by Chen et al. when DILIrank was assembled [13].

### 5.3 Featurization

For each compound we compute four descriptor blocks: (i) a 2,048-bit Morgan/ECFP4 fingerprint of radius 2 [53], (ii) the 167-bit MACCS structural key fingerprint [18], (iii) 15 physicochemical descriptors from RDKit [36] (topological polar surface area, octanol–water partition coefficient logP, hydrogen-bond acceptor/donor counts, total ring and aromatic ring counts, fraction of sp^3^ atoms, formal-charge counts, etc.; the full list matches the DILIPredictor DESCS block [55]), and (iv) the full Mordred 2D descriptor block [45] of ∼1,600 features. We then apply the same feature subset that DILIPredictor selected via stability-selected variance pruning on the LivTox training corpus [55]: 318 Morgan bits, 89 MACCS bits, all 15 physchem descriptors, and 435 Mordred descriptors, totaling **857 dimensions**. A 5-dimensional one-hot *training source identity* channel is appended at training time, one bit for each of the five sources {3, 5, 6, 7, 8} defined in Table 7, and fixed at inference time to the bit corresponding to source 7 (preclinical hepatotoxicity), yielding a 862-d input vector. The five source identity bits are denoted src_3, src_5, src_6, src_7, src_8; their role is to let the classifier calibrate against the source-specific positive class prior (which spans 0.34–0.75 across the five sources, Table 7) rather than collapsing all sources to a uniform label.

### 5.4 Oakuloid classifier

#### 5.4.1 Architecture: stacked GBDT, fingerprint random forest, and proxy assay ensemble

Oakuloid is a stacked ensemble that combines three gradient boosted decision tree (GBDT) backbones, LightGBM [33], XGBoost [14], and CatBoost [52], with four fingerprint random forests (one each on the MACCS, Avalon, atom-pair, and topological-torsion fingerprints) [10] and four predicted LivTox proxy-DILI features, all combined by a logistic regression meta learner. At inference, the model is evaluated as a single function: given a SMILES string, the feature vector is computed once (Section 5.3) and routed in parallel through the base learners producing base scores (out-of-fold scores are used only to train the meta learner) 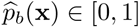 for each GBDT backbone *b* ∈ {lgb, xgb, cat}, the four fingerprint random forests, and the four proxy-DILI features, which the meta learner combines to a single hepatotoxicity probability

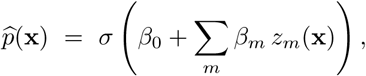

where *σ* is the sigmoid, *z*_m_ ranges over the GBDT base probabilities, the fingerprint random forest out of fold probabilities, and the proxy assay features, and the coefficients *β* are learned by L2-regularized logistic regression on out of fold predictions over the training set. In a separate ablation, expanding the predicted proxy assay panel from four to ten features contributed only +0.006 AUROC on the full TDC benchmark and +0.002 on the leakage clean subset; we therefore retain the four feature LivTox panel as the primary model and report the ten feature panel as a robustness check. Each base learner is class balanced (`class_weight=’balanced’` for LightGBM; `scale_pos_weight` = (1 − *ȳ*)*/*ȳ for XGBoost and CatBoost) and minimizes the binary cross-entropy

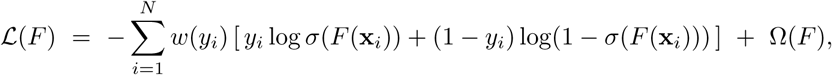

with *w*(*y*) = *N/*(2 |{*i* : *y*_i_ = *y*}|) the inverse-frequency class weight and Ω(*F*) the backbone-specific complexity regularizer.

#### 5.4.2 Model selection was restricted to groupwise cross-validation on the training set only

CatBoost hyperparameters were tuned by an Optuna search [2] whose objective is groupwise (InChIKey-14) cross validated AUROC on the training set alone. LightGBM, XGBoost and the fingerprint random forests use prespecified class balanced configurations selected before evaluation; evaluation labels and outcomes are never inspected during tuning or model selection. Evaluation compound identifiers are used only for the overlap-removal audit described above. Grouping folds by InChIKey-14 prevents stereoisomers and salts of the same compound from straddling the train/validation split, which would otherwise leak near duplicate chemistry into the validation score. We verified that this clean protocol generalises better than a test-set-in-the-loop alternative: a diagnostic probe that instead tuned directly on the TDC benchmark scored 0.581 on a fully held out novel chemistry slice (Geci), whereas the groupwise CV diagnostic model scored 0.613 on the same slice. This CatBoost-only diagnostic differs from the shipped full stack, whose doubly disjoint Geci AUROC is 0.590 (Supplementary Table S1). The resulting configurations are summarized in Table 8; the three backbones supply decorrelated decision boundaries that the meta learner combines (Section 2.1).

##### Why a stack instead of a single model

No single backbone is uniformly optimal across the internal and external cohorts. The stack is therefore evaluated as a held out prediction rule rather than as a training set fit: the logistic meta learner is trained only on groupwise out of fold base scores plus the four proxy-DILI features, and the final AUROC values are computed on datasets that are never used for tuning or meta learner fitting.

#### 5.4.3 Variance estimation

For each evaluation set, the entire stacked ensemble is retrained across five random seeds *s* ∈ {0, 1, 2, 3, 4} (each seed used uniformly for fold splits, base-learner internal RNGs, and meta learner training) and we report the five-seed mean ± standard deviation of the per-seed AUROC (TDC 0.809 ± 0.001; DILIrank 0.844 ± 0.010). Each seed re-randomizes the cross-validation fold splits, the base-learner internal RNGs, and meta-learner training. The released ensemble predictions used for the headline benchmarks reach AUROC 0.811 (TDC) and 0.850 (internal DILIrank): these are the deployed seed-ensemble (averaging the five seeds’ probabilities), reported alongside the per-seed means above rather than in place of them. Implementation uses LightGBM 4.6, XGBoost 3.2, CatBoost 1.2, and scikit-learn random forest and logistic regression components inside scikit-learn 1.8 [50] pipelines.

### 5.5 Compared baselines

We evaluate 13 prediction scores in total: Oakuloid structure, Oakuloid + BIO rank mean fusion, and 11 baseline scores from five method families. Two baseline scores are retained only as leakage positive overlap controls in the relevant benchmarks. The baseline families are: (i) wet lab IC_50_/C_max_ from the iBAC platform (Section 5.2); (ii) the upstream DILIPredictor (Seal 2024) [55] reporting three OOD-valid hepatotoxicity endpoints (DILIPred-Preclin, DILIPred-HumanHep, DILIPred-Mitotox) and one DILIst-FDA overlap-control endpoint (DILIPred-DILIst-FDA); (iii) ADMET-AI’s Chemprop directed message-passing neural network (D-MPNN) [76] ensemble [59] reporting DILI, Tox-Consensus, stress response mitochondrial-membrane-potential (SR-MMP), and stress response p53 (SR-p53); (iv) MolFormer-XL frozen mean-pooled embeddings (768-d) [54] paired with a LivTox-trained LightGBM head; (v) ChemBERTa-2 frozen embeddings (384-d) [1] paired with a LivTox-trained logistic regression head.

Where threshold-dependent metrics are reported, every method is scored at its own cohort Youden threshold. The native cutoffs of the third-party tools (≥ 0.5 for DILIPred; per-endpoint defaults for ADMET-AI) are used only for the clinical-stage binary flags (Table 2). BIO, Oakuloid, MolFormer-XL and ChemBERTa-2, whose heads were trained by us on LivTox, instead use cohort Youden thresholds. AUROC and three-level Spearman *ρ* are threshold free and directly comparable across all rows.

### 5.6 Statistical metrics

The threshold free primary metric is AUROC [25]. Threshold dependent secondary metrics (accuracy, F1 score (the harmonic mean of precision and recall), sensitivity, specificity, precision, and balanced accuracy) are computed at the Youden optimal threshold [77]. Unless otherwise stated, the Youden threshold of each model is derived on the same cohort on which the metrics are reported (the small-*N* DILIrank cohort or the strict external TDC subset); this convention biases threshold dependent metrics optimistically relative to a held out threshold protocol, and the threshold dependent values should therefore be read as operating point characteristics under each tool’s best-case threshold rather than as held out generalisation estimates. As a leakage-safe check on this optimism, we also recompute MCC under a held out threshold, with each method’s Youden threshold fitted on one cohort and applied to the other: Oakuloid’s operating point advantage persists on the strict external TDC subset (MCC 0.485 vs. DILIPred-Preclin 0.429) but reverses on the small internal DILIrank cohort (0.409 vs. 0.483), confirming that the cohort optimal MCC gap is an operating point characteristic rather than a comparative claim. AUROC remains the primary head-to-head metric throughout. Differences between paired AUROCs are tested with DeLong’s test [17] as the primary significance test for correlated ROC curves. We additionally report paired bootstrap 95% confidence intervals for AUROC differences (5,000 resamples) [19] for the key contrasts (Supplementary Table S12). We also report the three-level Spearman correlation [57] (*ρ*_3-level_) between the continuous score and the ordinal label (vMost = 2, vLess = 1, vNo = 0).

### 5.7 Wet lab fusion

We fuse the structural prediction with the experimental IC_50_/C_max_ ratio using *rank mean averaging*. For a compound with structural score *s* ∈ [0, 1] and wet lab ratio *r >* 0 the fused score is

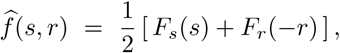

identical to the rank mean fusion equation stated in Section 2.3. The two natural choices for the empirical CDFs *F*_s_*, F*_r_ are: (a) within-cohort percentile ranks, *F*_s_(*s*_i_) = *rank*_pct_({*s*_j_}^N^_j=1_)_i_ and likewise for *F*_r_, with ties resolved by averaging, used for the benchmarking results in Section 2.3; and (b) fixed CDFs precomputed from a reference set (the 4, 681-compound LivTox training corpus for *F*_s_, the 122-compound DILIrank wet lab cohort for *F*_r_), released alongside the model and used for single-compound inference. The fusion has three useful properties: (i) it is invariant to monotonic rescaling of either input (the score *s* may be a probability or a logit, *r* may be in *µ*M or unitless, with the same fused output); (ii) it is robust to the heavy-tailed distribution of *r*, which spans ∼4 orders of magnitude in our cohort (0.02 ≤ *r* ≤ 1, 000); and (iii) when *r*_i_ is missing for some compound, we substitute the median rank *F*_r_(−*r*) = 0.5, which shrinks the fused score toward the structural-rank component; this imputation applies only when computing a fused score, whereas the deployed default for a compound with no measured ratio is the structural score alone (Figure 1). The equal-weight choice *w* = 1*/*2 is recommended throughout this work; it can be tuned to favour wet lab over structure when relative confidence is known (see Limitations).

### 5.8 Concentration analysis

We characterize the recoverable signal as a function of the assay concentration *C*^⋆^ = *k* × *C*_max_ for *k* ∈ {1, 3, 10, 30, 100}. For each compound *c* in the 132-cohort with *b*_c_ full-ladder plate batches and 3 technical replicates per (concentration, batch) cell, we draw IR(*C*^⋆^*, c*) as follows (Algorithm 9).

We additionally report the 3-level Spearman correlation [57] of the same IR(*C*^⋆^) score against the ordinal label {0, 1, 2} for vNo/vLess/vMost. Trial-to-trial scatter quantifies sensitivity to batch and replicate sampling; AUROC differences of *<* 0.01 between *ϕ* strategies indicate a noise floor below the across-batch signal.

### 5.9 Leakage audit

Following best practice for ML benchmarking on small chemistry datasets [32, 60, 67], we performed two complementary checks. First, we removed any TDC DILI compound whose 14-character InChIKey [26] matched either the Oakuloid training set (33*/*475 compounds removed, 6.9%) or the internal 122-compound cohort (41*/*475 removed, 8.6%); the two InChIKey-14 removal sets are disjoint (33 + 41 = 74 removed, 401 remain), and the resulting strict external subset of 401 compounds is used throughout the OOD evaluation in Section 2.1. Second, on the internal cohort we computed Morgan-2048 Tanimoto nearest neighbour distances [53] between each evaluation compound and the LivTox training set, and recomputed AUROC after progressively dropping near duplicates at thresholds *T* ≥ 0.85 and *T* ≥ 0.70 (Table 10), rescoring every method on the surviving compounds. Removing the closest neighbours left Oakuloid’s internal AUROC essentially unchanged (0.850 → 0.840) and did not reduce DILIPred-Preclin either (0.832 → 0.860), indicating that the internal-cohort AUROC is not driven by near-duplicate chemistry. The within-TDC similarity stratification (Section 2.2) separately shows that the comparative lead narrows on structurally novel compounds.

**Table 9:**
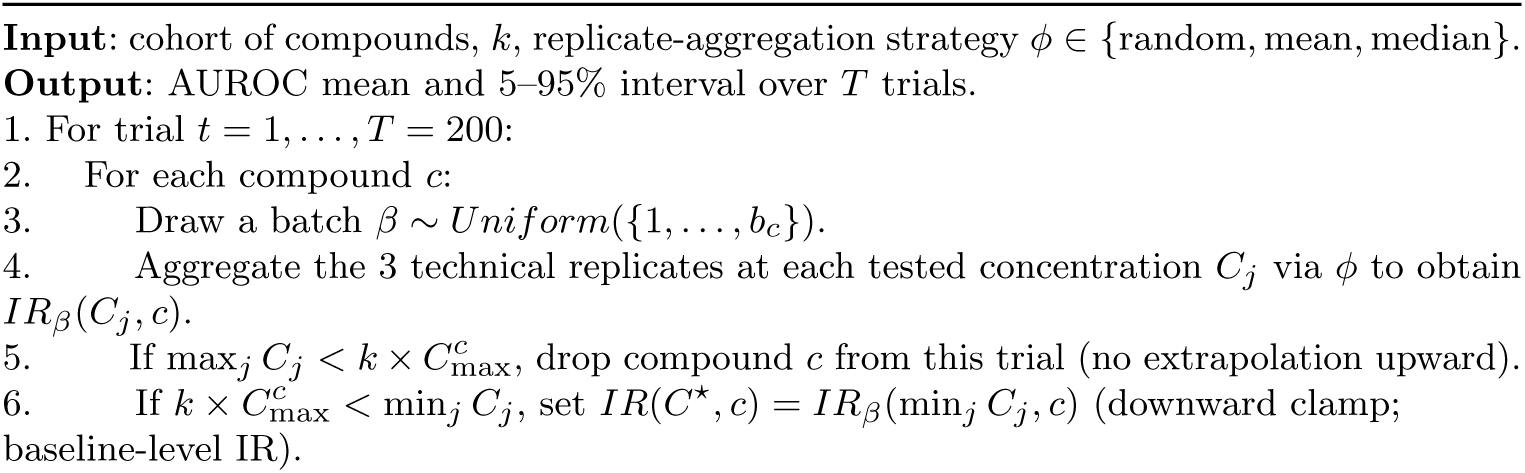
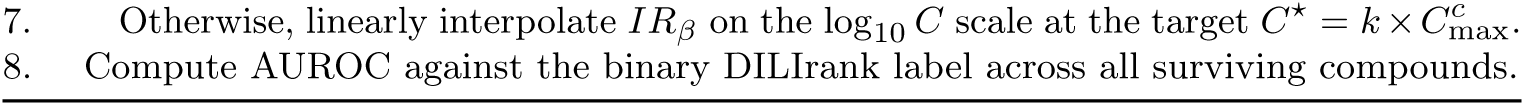
Algorithm 1: Monte-Carlo evaluation of single-point IR @ *k* × *C*_max_.

**Table 10:**
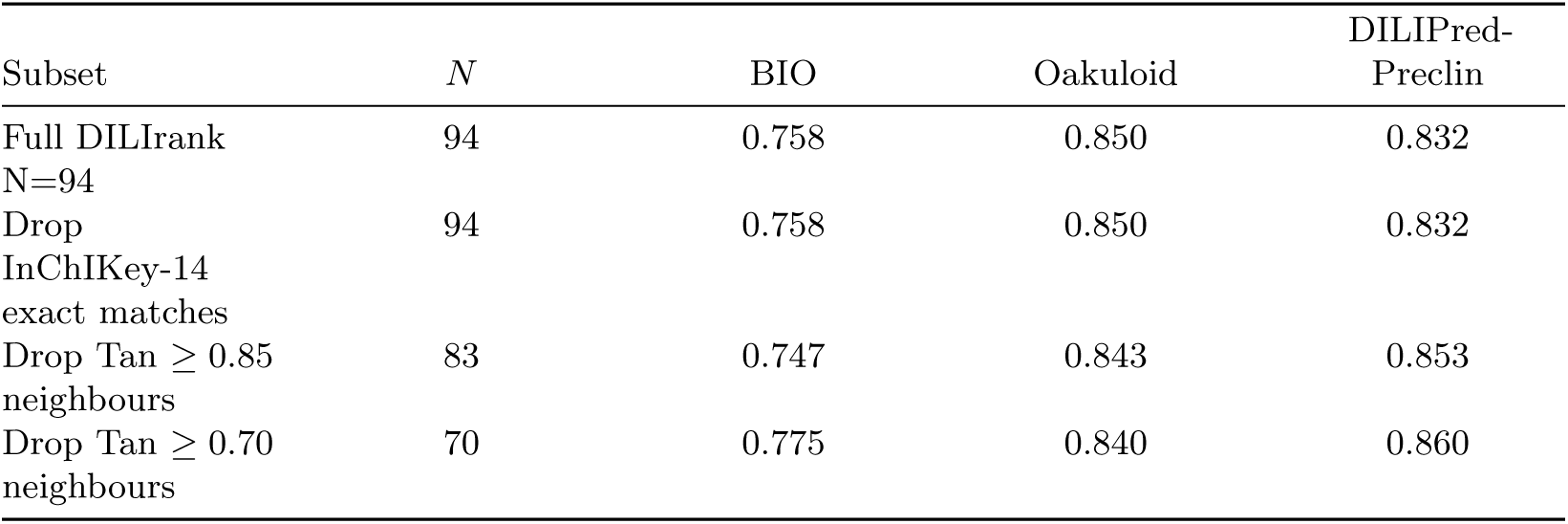
Tanimoto-stratified sensitivity analysis of the internal AUROC on the DILIrank binary cohort. Removing nearest training set neighbours does not degrade performance, indicating that the observed AUROC is not driven by training–test similarity.

Models whose training set InChIKey-14 overlap with the evaluation cohort exceeded 30% were excluded from the head-to-head comparisons. DILIPred-DILIst-FDA [61] is retained only as a training-overlap control. The full audit, including per compound exact-match flags and Tanimoto-nearest neighbour distances, is released as Supplementary Data 1.

Two provenance-specific safeguards complete the audit. First, the independent Geci set is scored by the structural model only. Because its DILI label is a deterministic threshold of the exposure-adjusted point-of-departure ratio (Section 5.1.6), any score that ingested that ratio would attain AUROC = 1.0 by construction. The exposure channel is therefore never fused into a Geci prediction and only structural model outputs enter the Geci columns. Second, within TDC we stratify by maximum Tanimoto similarity to the training set; as reported in Section 2.2, the benchmark lead is confined to the training-similar regime and is absent on the 24% of compounds with similarity *<* 0.4, the within-benchmark counterpart of the conservative Geci result.

Finally, a label-permutation (Y-scrambling) null confirms that the featurization carries nonrandom signal rather than a protocol artefact: retraining the CatBoost backbone on DILI labels shuffled within the training set (GroupKFold InChIKey-14 folds intact, three permutation seeds) collapses performance to chance, with AUROC 0.508 ± 0.038 on TDC-401 and 0.498 ± 0.011 on the internal out-of-fold predictions, versus 0.805/0.837 unpermuted.

## 6 List of abbreviations

ALT: alanine aminotransferase
ADMET: absorption, distribution, metabolism, excretion, toxicity
AUROC: area under the receiver operating characteristic curve
BalAcc: balanced accuracy
BIO: IC_50_/C_max_ wet lab score
BSEP: bile salt export pump
C_max_: maximum plasma concentration
CYP: cytochrome P450
DILI: drug-induced liver injury
DILIrank: FDA reference list of clinical DILI risk
D-MPNN: directed message-passing neural network
ECE: expected calibration error
F1: harmonic mean of precision and recall
FDA: U.S. Food and Drug Administration
GBDT: gradient boosted decision tree
iBAC: integrated biomimetic array chip
IC_50_: half-maximal inhibitory concentration
InChIKey-14: first 14 characters of the IUPAC International Chemical Identifier hash
IR: inhibition rate
LightGBM: light gradient-boosting machine
LivTox: LivTox proxy-DILI compendium
MACCS: molecular access system
MCE: maximum calibration error
OOD: out-of-distribution
PHH: primary human hepatocyte
Prec: pre-cision
QSAR: quantitative structure-activity relationship
RDKit: RDKit cheminformatics toolkit
ROC: receiver operating characteristic
SHAP: Shapley additive explanations
SMILES: simplified molecular input line entry system
Sens: sensitivity
Spec: specificity
SR-MMP: stress response mitochondrial mem-brane potential
SR-p53: stress response p53
TDC: Therapeutics Data Commons
TPR/FPR: true/false positive rate
vDILI-Concern: FDA DILIrank risk label {vMost, vLess, vNo, Ambiguous}
XGBoost: extreme gradient boosting.

## 7 Declarations

### 7.1 Ethics approval and consent to participate

Not applicable.

### 7.2 Consent for publication

Not applicable.

### 7.3 Availability of data and materials

The Oakuloid package, the primary 4-proxy model-training and inference pipeline, trained-model artefacts, and all manuscript figure-generation scripts are hosted at the project repository [79] under Apache License 2.0. Software information: project name, Oakuloid; project home page, https://github.com/DaXiangBio/Oakuloid; archived version, release v0.3.1 in the project repository; operating system, platform independent; programming language, Python; license, Apache License 2.0; restrictions on use, none. The accompanying datasets are released as Supplementary Data files: per compound leakage audit (Supplementary Data 1), full 862 feature SHAP and source identity audit (Supplementary Data 2), the 122-compound DILIrank wet lab dataset with SMILES (Supplementary Data 3; also tabulated in Supplementary Table S11), per compound predictions across all 13 prediction scores tested in this work with binary calls (Supplementary Data 4), and the 132-compound dose-response raw-IR readout (Supplementary Data 5, 1,909 rows). External datasets used: DILIrank vDILI-Concern labels [13], the LivTox proxy-DILI compendium [55], TDC DILI [28], and the Geci et al. independent hepatotoxicity dataset [21]. In line with FAIR data principles [71], a persistent archive (Zenodo DOI) snapshotting the exact code, the five Supplementary Data files and the trained-model artefacts will be deposited on acceptance, so the analysis is reproducible independently of the mutable repository tag.

### 7.6 Authors’ contributions

Fan Zhang and Yu Zhou contributed equally to this work. Yu Zhou conceived the core methodological ideas and designed the clean-CV stacked ensemble, fingerprint random forest, and proxy feature framework of Oakuloid. Fan Zhang implemented Oakuloid, ran all benchmarking and ablation experiments, prepared the figures, and wrote the initial manuscript. Duanchen Ding and Feng Zhang performed the experimental IC_50_/C_max_ measurements on the iBAC 3D-PHH platform and curated the C_max_ master table. Rong-Rong Xiao and Xiaoni Ai conceived the study, supervised the project, acquired funding, and revised the manuscript. All authors read and approved the final manuscript.

## 7.7 Acknowledgements

We thank Srijit Seal and the DILIPredictor authors for releasing the LivTox proxy-DILI compendium under MIT license; reused components are attributed in the project NOTICE file as required by Apache License 2.0. We thank IBM Research for releasing MolFormer-XL weights and DeepChem for ChemBERTa-2. We acknowledge the iBAC 3D-PHH platform team: Tian Lv, Xia Tu, Peiwen Li, Tiantian Wang, Haiheng Dong, and Pengfei Tu, whose work establishing the wet lab assay [73] provided the IC_50_/C_max_ measurements central to this study. Fan Zhang used Anthropic’s Claude (Opus 4.7) coding assistant during literature search, code refactoring, and editorial drafting; all scientific claims, statistical analyses, and final manuscript text were verified, edited, and approved by the human authors. No content was generated and accepted unverified.

